# Efficient task generalization and humanlike face perception in models that learn to discriminate face geometry

**DOI:** 10.64898/2026.01.31.703048

**Authors:** Seojin Lee, Josh Ying, Ahana Dey, You-Nah Jeon, Elias B. Issa

## Abstract

Artificial deep neural networks (DNNs) can excel at face recognition from 2D photographs where both shape and appearance cues abound; however, DNNs have rarely been challenged to recognize faces strictly based on face geometry. Here, we show that DNNs, even those fine-tuned on face photographs, had almost no generalization performance to a new geometry-based face task, while in the opposite direction, networks fine-tuned only on geometrically defined, textureless faces readily generalized to textured faces. To learn geometry in a more practical setting with colored and textured faces, we trained discrimination on face emotion in addition to face identity, which resulted in less texture bias and generalized well across face tasks. Learning in this way from just four individuals and their expressions generalized to unseen individuals, even exceeding standard models which are trained on classifying hundreds of face identities. Compared to standard models, emotion and identity trained models developed more humanlike errors in the identities or emotions that they confused. This novel method learns in a humanlike manner using only a few individuals but enriched with expressions that widely vary face geometry – similar to early human experience during child-parent interactions. Thus, this bioinspired work has broad implications for how moving toward humanlike learning of geometry in artificial vision can be both highly sample efficient and highly performing.

## Introduction

Faces pose a unique challenge for both human and machine vision because a vast number of identities must be discriminated using only subtle variations within a common facial structure^1^. Because face identity and expression vary through fine shape changes within a generally preserved configuration, face recognition places stronger demands on visual discrimination than general object recognition. Yet humans excel at this task, achieving rapid and reliable identification whilst also discerning their gaze and expression across wide variations in face pose and lighting^2,3^. Remarkably, even when face images are severely degraded, face recognition remains robust, failing only at very low resolutions^4,5^. Thus, face recognition represents a final challenging frontier for artificial vision systems to truly master.

Recent advances in artificial deep neural networks (DNNs) have led to breakthroughs in face recognition, with models such as CASIA-WebFace^6^, FaceNet^7^, and VGGFace2^8^ achieving state-of-the-art performance on real world face images. Beyond performance-driven DNNs, recent computational approaches have aimed to explain how human face recognition may work by using more explicitly face specific algorithms. Active appearance models describe faces using joint statistical variation in shape and texture^9–11^, while the inverse graphics approaches use probabilistic generative models to infer latent 3D factors such as identity, pose, and expression^12,13^. Recently, DNNs have also been used to probe human face perception, revealing that human behavioral signatures emerge in face-trained networks and that training objectives lead to alignment with neural data^13–16^.

Despite these recent successes, our behavioral work on humans, monkeys, and DNNs showed that DNNs that previously excelled at face and object recognition in natural images were unable to perform a fundamental geometry-based face recognition task (GFR)^17^. In contrast, both monkeys and humans performed GFR at a similarly high level to their object recognition performance, showing no comparable drop in accuracy in primate vision. This work exposed a face-specific gap in current discriminatively trained DNNs, highlighting a fundamental weakness in how these models represent facial geometry.

In diagnosing this failure of standard DNN training, one possibility is that overlying color and texture cues present in natural faces may subvert any true learning of underlying geometry. In such a scenario, models can achieve high face recognition performance by using relatively view-invariant texture cues – such as hair color or skin tone – without learning the geometric structure that supports robust generalization across appearances. An alternative to such large-scale face training with thousands of identities is the hypothesis that face learning actually proceeds by exposure to only a few, socially important faces that a child frequently interacts with on a daily basis, with broader refinement occurring later as social experience expands. At first, this biologically grounded yet sample-efficient strategy might appear inherently limited, as training on only a few individuals could lead to severe overfitting to their specific features, hence undertraining for the larger face domain. On the other hand, there is a rich, high-dimensional training diet offered by a single face when presented under different viewpoints, lightings, and expressions – information that is likely leveraged by biological vision. Here, we explored whether there were computational benefits to this alternative face-training regime and how best to leverage it to remedy the lack of generalization performance and task transfer observed under standard large-scale identity training.

We developed a synthetic face stimulus framework for exerting direct control over native 3D facial mesh geometry manipulated through structured identity or expression changes, independently of texture. Using this framework, we found that infusing emotion recognition into model objectives not only biased learned features toward geometry-based, rather than texture-based features, it led to efficient generalization across many face tasks – transferring to both identity and emotion recognition, with and without texture cues. This learning benefit was evident for both supervised and self-supervised pretrained backbones, suggesting the general applicability of this new training approach. Finally, we asked whether the final, learned representations produced behaviors that resembled those of face recognition behavior by humans. Remarkably, the particular identity and emotion confusions humans made were produced by our sample efficient face learning method but not by standard training methods. Together, these results suggest that building in more geometry based strategies to discriminate faces may prove essential to matching biological vision in its learning efficiency, generalization performance, and behavioral idiosyncrasies.

## Results

### Standard DNNs failed to generalize to geometric face recognition tasks

Feedforward deep convolutional neural networks (DNNs) are known to exhibit a texture bias – a tendency to rely more on surface appearance than on shape. We hypothesized that such texture-reliant representations would struggle on tasks that required identity discrimination based solely on 3D face shape. To test this, we curated a geometry-based face recognition task (GFR) which required a binary distinction between two identities that only differ in face geometry, with all texture and color information removed (**Fig. 1A**). Faces, which were always centered in the image, varied across identity-preserving transformations including 3D viewpoint, scale, lighting direction, and background. We evaluated whether standard, off-the-shelf DNNs trained for object or face recognition could generalize to geometry-based face discrimination. Specifically, we tested ImageNet-supervised ResNet-50^18^, AlexNet^19^, and VGG-16^20^, as well as face-specialized models including VGG-Face^8^ and FaceNet (128-dimension and 256-dimension embeddings)^7^. The full battery of GFR evaluation included 66 binary classification tasks (2AFC), corresponding to all unique pairs among 12 individual identities collected in the lab (**Fig. 1B**). For each model, we extracted penultimate-layer features and trained a linear Support Vector Machine for each identity pair; overall GFR accuracy was computed as the mean accuracy across all 66 face pairs.

**Figure 1.**
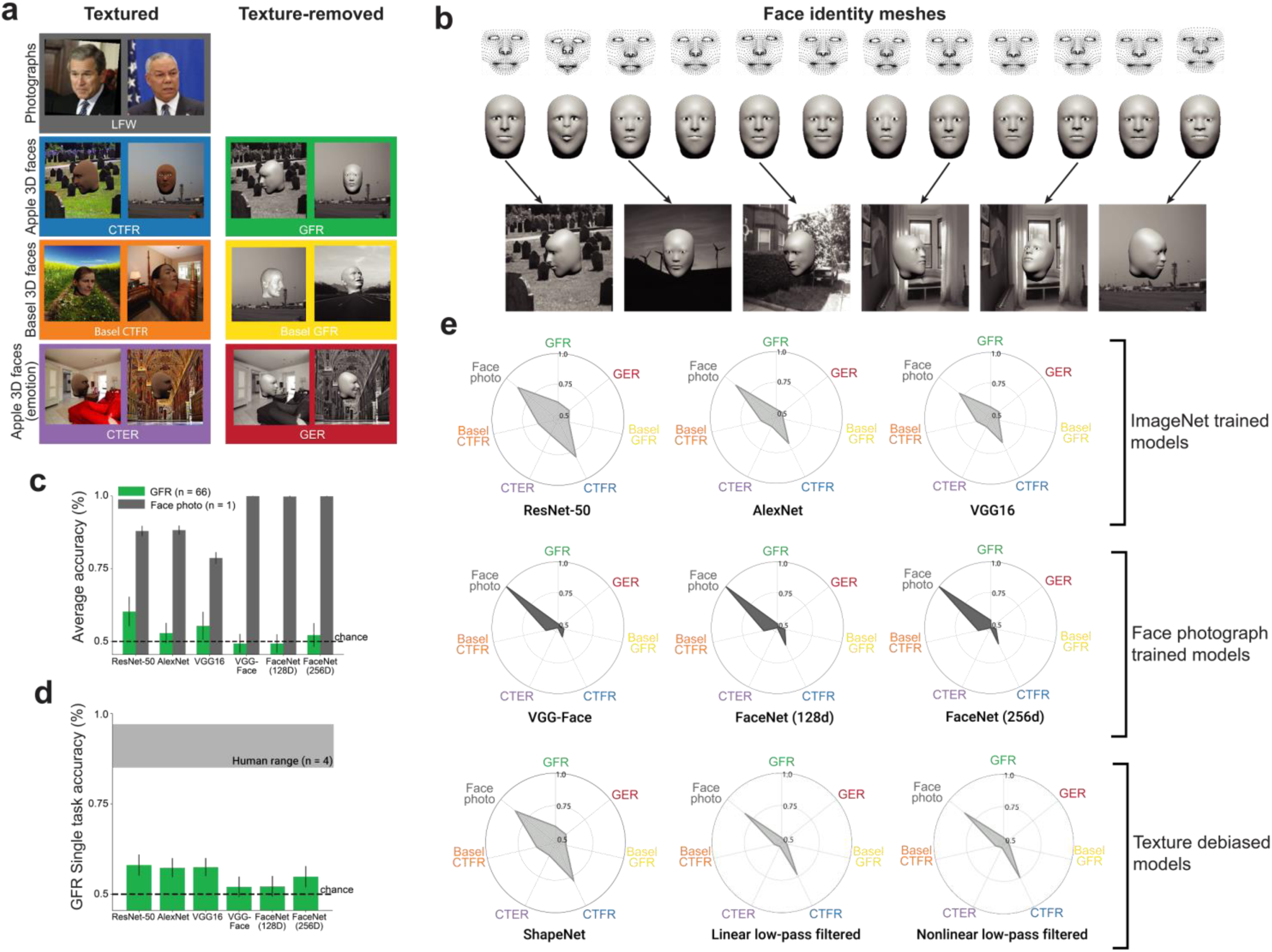
Standard models pretrained on objects or faces performed poorly on geometry based face tasks. **a,** Example stimuli from seven face recognition tasks varying in stimulus type (photographic versus 3D-rendered), recognition goal (identity versus emotion), and the presence of texture. Left, textured versions with naturalistic color, texture, and backgrounds. Right, texture-removed faces with grayscaled backgrounds. **b,** Generation of 3D face stimuli from mesh data, varying only internal face geometry across individuals while matching external head geometry, eye, color, and texture cues. Each column shows one identity. Top, raw 3D geometry defined by 936 vertices across key facial regions. Middle, same geometries stitched onto a standardized 3D head model with identical black spheres used as pupils, then rendered as a smooth, textureless monochrome surface with lighting. Bottom, example training images, generated by rendering face meshes under a random rotation, scaling, lighting variation and background composition. **c,** Model accuracy shows a clear gap between the photograph-based (Face Photo) and geometry-based (GFR) face identification tasks. Each GFR bar shows mean binary task accuracy across the 66 identity pairs, and the Face Photo task utilized one pair from Labeled Faces in the Wild. **d,** DNN performance fell well below human performance on the binary task, which was measured on a single face pair (gray shaded region: human = 88–97)^17^. **e,** Polar radar plots summarizing DNN performance across the seven face tasks shown in panel ***a***, arrayed radially, with normalized accuracy indicated by radial distance. Larger areas circumscribed by the performance polygon indicate greater generalization across tasks. All standard, off-the-shelf models displayed relatively narrow performance profiles, indicating limited generalization beyond identification in standard face photographs. Notably, face-trained models (VGG-Face, FaceNet-128d, FaceNet-256d; middle row) showed the narrowest profiles, performing near ceiling on Face Photo while failing elsewhere. Even models with reduced texture bias by training on Stylized ImageNet or low pass filtered images still largely failed to generalize outside of face photographs to textureless geometry-based face recognition (bottom row).

Despite their strong performance on conventional face benchmarks, all tested DNNs failed to generalize to the GFR task. ResNet-50 achieved 60.3% accuracy, only slightly above chance, and the remaining models performed near chance levels, even those that were specifically trained on faces such as VGG Face or FaceNet (**Fig. 1C**). These results are consistent with the hypothesis that both object-trained and face-specialized DNNs are biased toward texture, extending prior work on texture bias in general object recognition^21,22^.

To confirm that this failure was specific to geometry-based generalization rather than a general deficit in face discrimination, we evaluated the same models on a standard binary identity classification task using images of two famous individuals (George Bush and Colin Powell) from the Labeled Faces in the Wild (LFW) dataset (**Fig. 1A**). On this benchmark, both face-specialized and object-trained models performed substantially better than on GFR (**Fig. 1C**). Face-specialized models achieved near-perfect accuracy, while object-trained models also reached high performance, though with greater variability. These results confirmed that the poor performance on the GFR task was not due to a general inability to discriminate faces. Instead, it highlighted the specific challenge posed by geometry-based face recognition, which requires identity discrimination from shape alone while remaining invariant to changes in 3D viewpoint, scale, lighting direction, and background.

We then tested whether models explicitly designed to reduce texture reliance would show improved performance on geometry-based face recognition. Specifically, we tested Shape-ResNet, which is trained jointly on Stylized ImageNet and ImageNet with additional fine-tuning^22^, as well as two models trained on low-pass filtered images using either linear or nonlinear blurring^23^. None of these models showed improvement on GFR, with all variants operating at or near chance levels (**Fig. 1E**, bottom row). These findings suggest that existing domain-general strategies for mitigating texture bias are insufficient for geometric face recognition.

The limitation of standard DNNs in the GFR task domain stood in stark contrast to biological visual capability where humans performed GFR effectively as well as they performed object recognition as shown in our recent work^17^. On a representative identity pair that was psychophysically tested, human participants performed near ceiling (**Fig. 1D**). In contrast, all DNNs tested on the same task performed near chance across architectures, underscoring the gap between artificial and biological vision in sensitivity to facial geometry.

We next asked whether the poor performance observed on geometry-based face recognition could be recovered by restoring facial texture and re-measuring performance on the same faces. In this colored, textured face recognition task (CTFR), models were evaluated on the same 66 identity pairs as the GFR task but with identity-specific facial texture overlaid onto the underlying 3D geometry (**Fig. 1A**). While faces still varied in viewpoint, lighting, scale, and background, the addition of texture selectively improved performance for some models, particularly for ImageNet pretrained models (ResNet-50: 80.22%, 13-20% performance gains), while face-specialized models showed only modest improvements (5–12%) despite extensive prior training on color-textured face datasets (**Fig. 1E**). This result underscores a critical limitation: although these models were optimized for face identity recognition, their reliance on texture learned under narrow training setting was insufficient to support generalization across novel textures and naturalistic variation in 3D viewpoint, lighting direction, and scale. This pattern was replicated using an independent set of face meshes and textures from the Basel Face Model^24^, where models again performed near chance when only geometry cues were available and showed only marginal gains when texture was included.

Given that these model performances on face identification reflected a reliance on face texture and an inability to extract face geometry information, we reasoned that models would generally fail on view invariant emotion recognition tasks. We therefore constructed two additional task batteries – geometry-based emotion recognition (GER) and color-texture-based emotion recognition (CTER) – that paralleled GFR and CTFR but probed emotional variation within a single identity (**Fig. 1A**, fourth row). Across all models, performance on GER (21 binary tasks across seven emotions) remained near chance, indicating that similar limitations observed for geometry based identity recognition extended to emotion recognition. Intuitively, adding texture may provide little diagnostic information in the domain of emotion recognition since texture remains effectively constant across expressions within an individual. Consistent with this intuition, texture addition did not improve performance on CTER and in some cases slightly reduced accuracy. Successful performance on both GER and CTER therefore required sensitivity to subtle, expression-driven geometric changes in facial features, such as mouth curvature and brow position. The complete pattern of performance for each model across identity and emotion recognition tasks, with and without texture, is summarized in the polar plots of **Fig. 1E**, which show a narrow band of high performance corresponding to a lack of generalization across face tasks.

### Training based on face geometry alone enabled broad generalization to both textured and textureless face tasks

To address the limitations observed in standard DNNs in recognizing faces based solely on their shape, we tested whether explicitly training a network on geometrically defined faces, where the face is stripped of any identifying texture or color cues, could enhance generalization even when appearance cues are later reintroduced. We fine-tuned an ImageNet-pretrained ResNet-50 model on an identity discrimination task using the same grayscale 3D face stimuli from our benchmark GFR task. During training, faces presented centrally varied across viewpoint, lighting, scale, and background, encouraging shape-sensitive representations invariant to these transformations and less reliant on surface-level appearance. We trained a set of models (GFR-trained models) using identity sets of 2, 4, 6, or 8 individuals with a fixed total number of training images at 100,000 per identity (see **Methods**). Following training, models were evaluated across seven face tasks spanning geometry- and texture-based domains.

GFR-trained models exhibited consistent performance gains across all tasks, with accuracy improving consistently as the number of training identities grew (**Fig. 2A,** left panel). On the in-distribution GFR task, these models substantially outperformed the baseline ResNet-50, with comparable improvements on out-of-distribution (OOD) meshes. Notably, rather than just hyperspecialize on the geometry only condition, these improvements extended to textured face recognition: on CTFR, GFR-trained models outperformed the baseline on CTFR (85.0% on OOD meshes versus 80.2%) despite never being exposed to textured faces during training.

**Figure 2.**
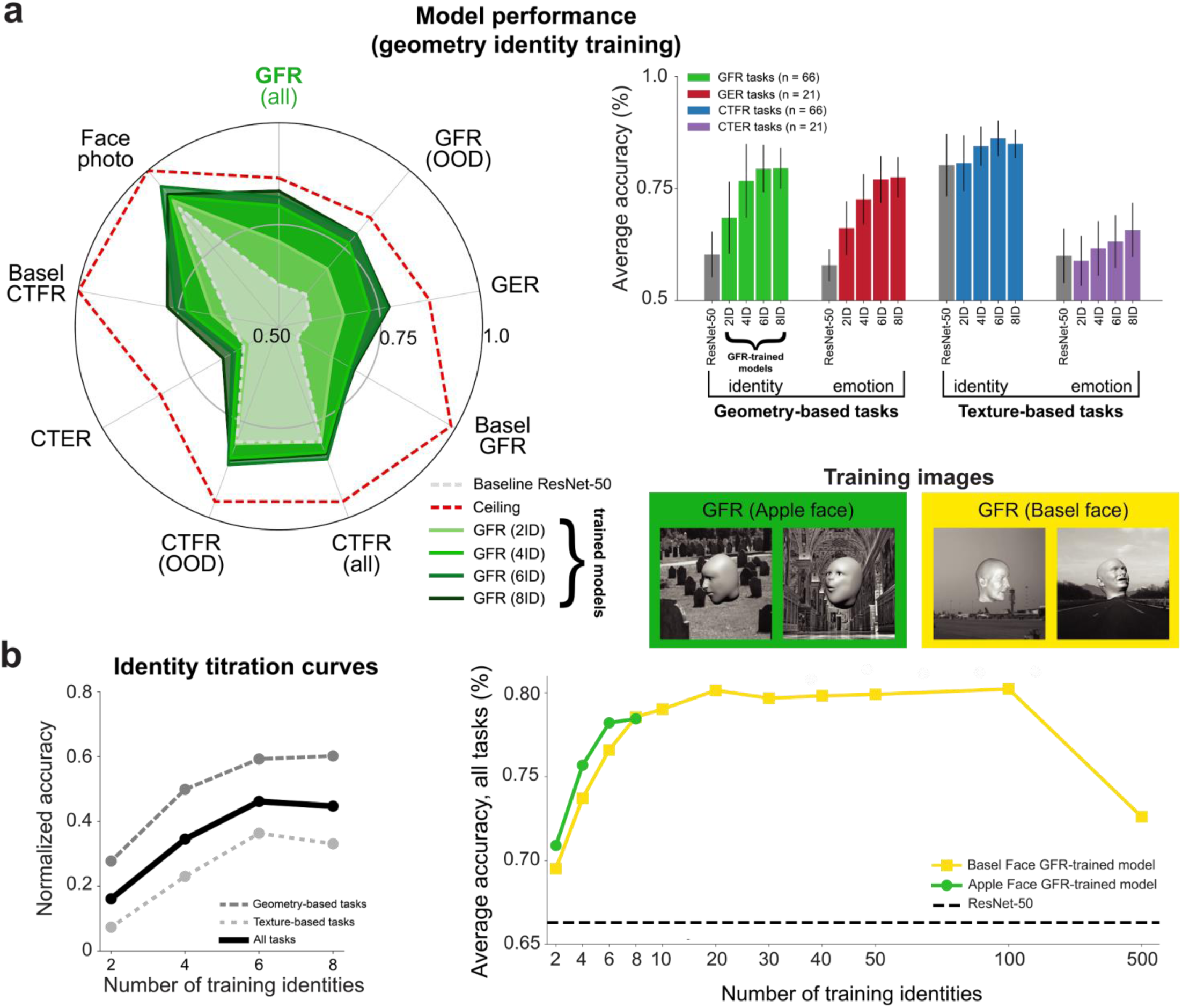
Geometry trained models exhibited performance gains across a broad array of face tasks. **a,** Generalization profile of GFR-trained models across task domains. Polar radar plot shows accuracy (radius) of models across the same nine evaluation tasks as in Fig. 1e. Green lines represent GFR-trained models with 2, 4, 6, or 8 training identities (darker shades indicate more identities). The dashed grey line marks performance of a baseline ResNet-50, and the red dashed line indicates the ceiling on each task (highest observed accuracy across all trained models, typically the model fine-tuned specifically on that task). GFR-trained models showed broad cross-domain generalization, with the largest gains on geometry-based tasks (GFR, GER, Basel GFR) and moderate improvements on texture-based tasks (right panel). **b,** Normalized accuracy as a function of the number of training identities for GFR-trained models using Apple face stimuli (left panel). Normalized accuracy for each task was scaled from the ResNet-50 baseline (floor) to the best-performing model (ceiling). Performance improved with identity diversity across all domains, but saturated around six identities. A similar finding was observed when training with the larger corpus of Basel faces where performance largely saturated early around ten identities (right panel, yellow curve).

Generalization also held across new stimulus domains. On Basel Face Model meshes – unseen during training – GFR-trained models generalized strongly, consistently outperforming the ResNet-50 baseline on both Basel GFR and Basel CTFER. Similar benefits were observed for emotion discrimination, where GFR-trained models far exceeded baseline performance on GER (66–77% vs. 58%) and matched baseline performance on CTER. These models also transferred well to real-world face photographs, achieving 90–95% accuracy on the Face Photo task. This generalization pattern of GFR-trained models stands in sharp contrast to the opposite direction of learning and transfer, where models trained directly on textured 2D face photographs failed to generalize to textureless face tasks (**Fig. 1E**). For example, the three face-specialized models, VGG-Face, FaceNet-128D and FaceNet-512D, achieved effectively chance performance on Apple Face GFR (49–52%), Apple Face GER (47-48%), and Basel GFR (51–52%) tasks.

To illustrate these generalization patterns, we visualized performance on four representative tasks – GFR, GER, CTFR, and CTER – as a function of the number of training identities (**Fig. 2A**, right panel). Performance gains scaled with training diversity and were strongest for geometry-based tasks, with smaller but reliable gains for texture-based tasks. Together, these results showed that training on facial geometry alone supported broad generalization across face recognition tasks without sacrificing performance in texture-rich domains.

### Geometry-based face learning performance saturated after a few identities

Having established that geometry-based training enables broad generalization across both geometry-and texture-based tasks, we next asked how many identities are required for this benefit to emerge. Standard face recognition models are typically trained on hundreds or thousands of identities^25,26^, but it remains unclear whether such large-scale identity diversity is necessary when training is grounded in 3D face shape rather than face appearance. By training GFR models with progressively larger numbers of identities and comparing cross-task performance, we found that normalized accuracy improved with increasing identity diversity but began to plateau after just six identities. Mean normalized accuracy across all seven evaluation tasks increased with the number of training identities and showed no further improvement beyond six identities (**Fig. 2B**, solid line in left panel). To examine this trend by task type, we grouped tasks into geometry-based (GFR, GER, Basel GFR) and texture-based (CTFR, CTER, Basel CTFR, Face Photo) categories. Normalized accuracy averaged across geometry-based tasks increased with identity diversity and saturated at six identities, with minimal gains thereafter (**Fig. 2B**, dark gray dashed line). Texture-based tasks showed a similar saturation pattern, improving up to six identities before declining slightly at eight identities (**Fig. 2B**, dark gray dashed line). Together, these results suggested that the generalization benefits of geometry-based training, as measured by normalized accuracy, saturate at approximately six identities regardless of task domain.

To test whether this saturation point holds in face training regimes drawing on larger face databases with potentially more features than our custom Apple Face stimuli, we turned to the Basel Face Model, which provides a substantially broader range of identity variation. Using this dataset, we trained Basel GFR models with 2 to 500 unique identities and evaluated their accuracy averaged across all Apple and Basel test tasks (**Fig. 2B**, right panel). Average accuracy increased sharply at low identity counts and reached a plateau at intermediate identity counts. Beyond this range, additional training identities yielded no further gains and eventually reduced performance. These findings confirmed that generalization effects of geometry-based training saturate at a relatively low number of training identities – around six when training using Apple Face stimuli, and under twenty with the larger Basel dataset. Importantly, this saturation occurred at an order of magnitude fewer identities than the hundreds typically used in large-scale supervised face recognition training.

### Training on textured faces does not generalize cross-domain to textureless faces

So far, we have shown that models trained to recognize identity from face geometry can generalize not only to other novel face geometries but also to textured synthetic faces and natural photographs. We next asked how good face generalization performance depended on the nature of the face training. As shown earlier in the polar radar plots, models trained conventionally on 2D textured face photographs failed to generalize across tasks, performing at chance level on nearly all evaluation tasks except Face Photo (**Fig. 1E**). This failure of conventional training on our wider face task battery could be simply driven by a lack of systematic scene variation during training that limited generalization across appearance conditions (e.g., face pose, lighting direction and background identity) that were part of our specific face stimuli – an issue that could be easily remedied by exposure to these variation conditions. Alternatively, the presence of identity-specific textures may lead models to overfit on surface-level appearance cues rather than extract more generalizable information about the underlying facial geometry, representing a more fundamental, inherent limitation.

To disentangle these possibilities, we created a color-textured face training set (CTFR) by overlaying identity-specific face textures onto our 3D shape-defined face meshes. This approach controlled geometric variation while introducing color and texture (**Fig. 3A**). CTFR-trained models were fine-tuned using the same identity sets and training protocols as GFR-trained models, allowing for direct comparison to the observed benefits of geometry-based face training. In stark contrast to their GFR-trained counterparts, CTFR-trained models revealed a marked asymmetry in generalization behavior, as they performed well on texture-based tasks but failed to generalize to geometry-based tasks. On the in-domain CTFR task, performance was substantially higher than the baseline (89-94% versus 80%), with similar advantages observed for held-out (OOD) textured meshes. These models also performed well on natural photographs, achieving 91–93% on the Face Photo task. However, performance on geometry-driven tasks showed no improvement: accuracy on GFR remained near the baseline models’ for both in-domain and OOD meshes, and similar limitations were observed for GER (**Fig. 3A**). Crucially, this limitation extended to CTER, where texture provided little diagnostic information about facial expression; hence, accuracy remained near baseline. The asymmetry between textured versus untextured faces for texture trained models persisted in the Basel Face dataset, as models performed poorly on Basel GFR but showed strong gains on Basel CTFR (**Fig. 3A**). Together, these results highlight a clear asymmetry in generalization: whereas geometry-based training transfers robustly across texture-rich and texture-free domains, texture-based identity training supports only within-domain improvements and fails on tasks requiring sensitivity to underlying 3D structure. Thus, even when we matched the training presentation context using our colored, textured faces – including variations in viewpoint, lighting, scale, background, and the style of rendering 3D faces into 2D images – it was the face’s texture itself similar to conventional face photographs that constrained task generalization after training with colored, textured faces.

**Figure 3.**
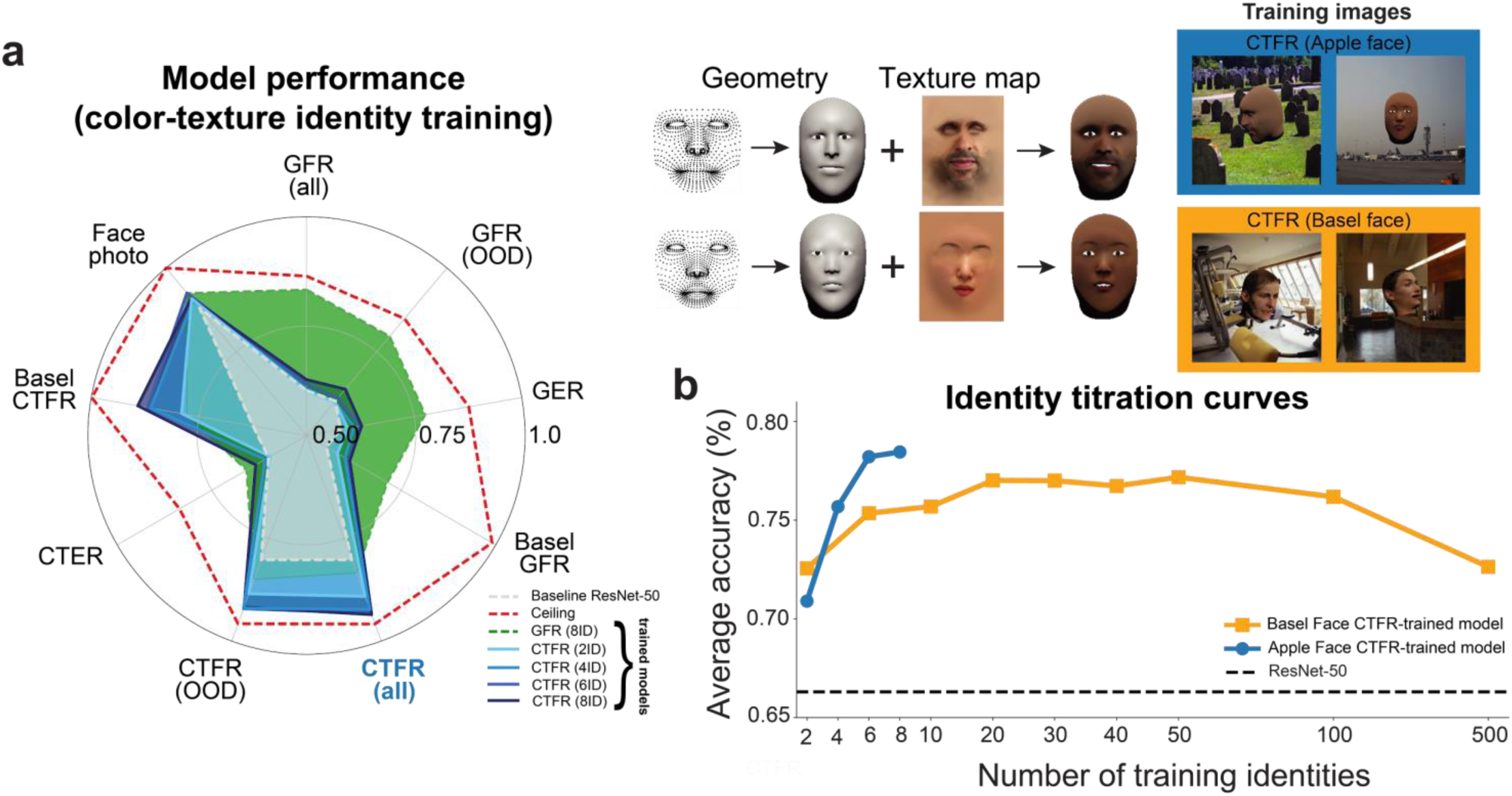
Models trained on colored-textured synthetic faces only generalized within colored-textured face tasks. **a,** Training models using colored-textured synthetic Apple faces did not generalize to geometry based tasks such as textureless face identification, nor to colored-textured emotion recognition (CTER) (blue regions in polar radar plot). For comparison, the geometry trained model (GFR-8ID, green region) did show broad performance gains (dashed gray and red lines indicate baseline ResNet-50 and ceiling performance, respectively). Example realistic textured face training images are shown on the right, generated by applying identity-specific texture maps derived from color photographs to the mesh surfaces. **b,** Average accuracy across all face tasks saturates under 20 training identities for both Apple face meshes and the larger corpus of Basel face meshes (same format as Fig. 2b; dashed black line indicates performance of baseline pretrained ResNet-50 model).

Importantly, simply increasing identity diversity could not resolve the limits on task generalization. Even when scaling to hundreds of identities in the larger Basel Face dataset, Basel CTFR-trained model performance continued to hover near the baseline ResNet-50 on geometry-based tasks (**Fig. 3B**). These results suggest that increased identity variation alone is insufficient to emphasize geometric variation during learning as it does not overcome the fundamental generalizability limits of texture-based face training.

### Training to perform emotion recognition in textured faces led to broad generalization performance

A key question is how to unlock the advantages of learning from face geometry without direct access to this information. Compared to the conventional approach of scaling up identity diversity alone in the hopes of capturing all features useful for robust face recognition, we explored a more efficient, biologically inspired learning regime motivated by early human development, where infants are repeatedly exposed to a small number of familiar faces expressing a wide range of emotions.

Simply training a model for emotion recognition using images from a single identity (with no identity labels, since only one individual was present), on its own produced general improvements spanning multiple face tasks. These gains increased with the inclusion of additional identities and peaked at four identities, where the model was trained on 28 categories formed by combining four identities with seven emotional expressions (neutral, happy, sad, anger, disgust, fear, surprise; **Fig. 4A**), yielding a 28-way classification task (CTFER-4IDx7EM). Though trained exclusively on colored, textured faces without direct access to shape information, the CTFER-4IDx7EM model stood out in generalizing to geometry based tasks, substantially outperforming both the ResNet-50 baseline and all CTFR-trained models (**Fig. 4A**). Notably, this higher level of performance was achieved using only four identities, whereas CTFR-trained models in the previous section showed little improvement over baseline even when trained on eight or more identities. At the same time, the model retained strong performance on texture-based tasks, well above baseline performance and comparable to even the best CTFR-trained models. Performance on the Face Photo task also improved to 93%, surpassing both the ResNet-50 baseline and all CTFR-trained models. Together, these results indicated that emotional variation provided an effective training signal for learning shape-sensitive representations, in spite of, and without interfering with, representation of textured faces for identity classification.

**Figure 4.**
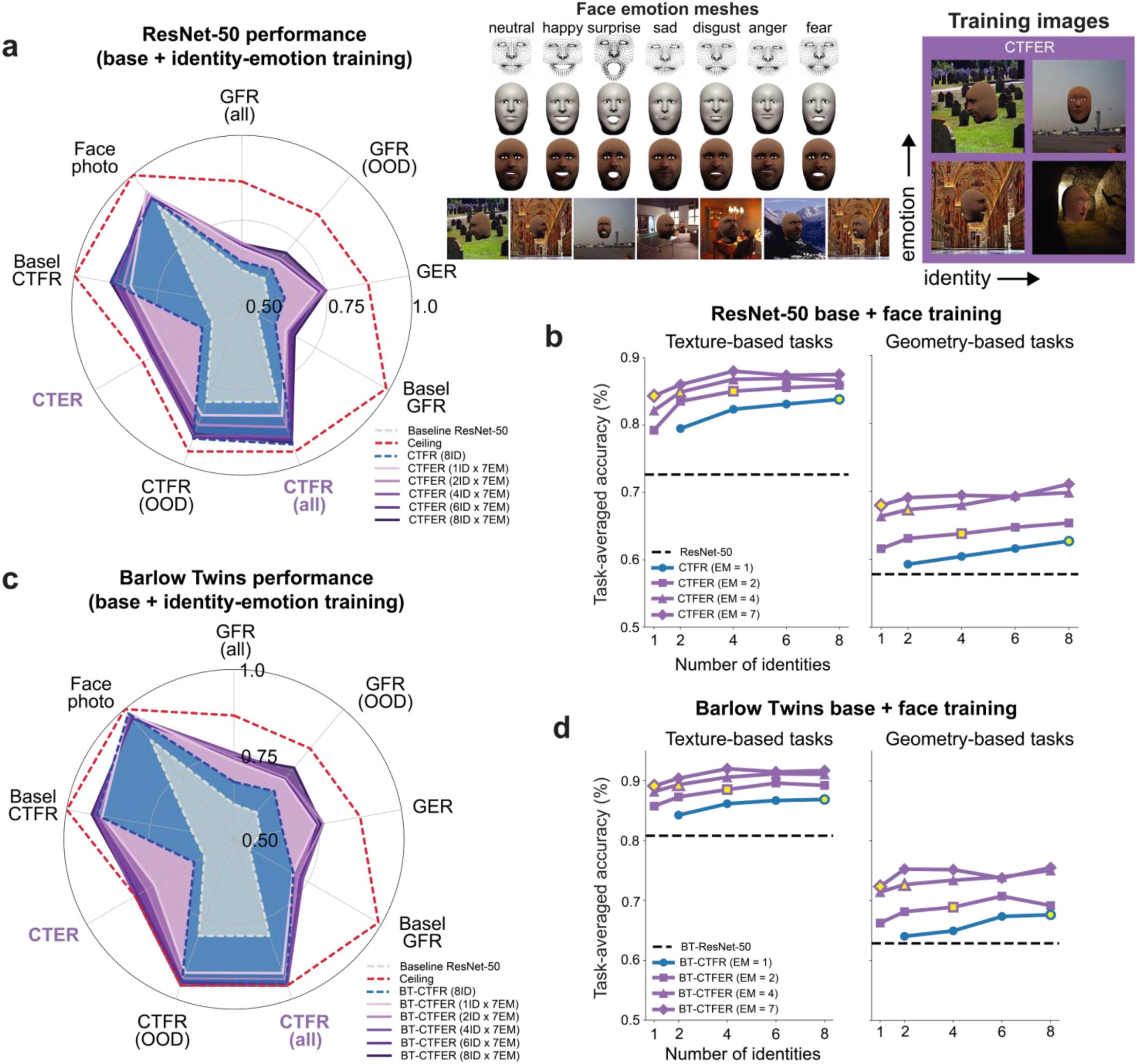
Models trained on both identity and emotion recognition from colored-textured faces exhibited broad generalization performance. **a,** Overall generalization profile across tasks, shown in the same format as Fig. 2a. Polar radar plots show performance for CTFER-trained models trained with 1, 2, 4, 6, or 8 identities, each combined with seven emotional variations (purple lines; darker shades indicate more training identities). Compared with strictly identity trained CTFR models which were relatively poor at geometry tasks (blue), identity and emotion trained CTFER models showed stronger and more balanced generalization across geometry- and texture-based tasks (purple). Shown at right, seven basic facial expressions applied to one identity to generate expressive, color-textured face stimuli for model fine-tuning and testing. **b,** Impact of increased emotional variation on generalization relative to simply increasing number of face identities. Left, task-averaged accuracy across geometry-based tasks for CTFR- and CTFER-trained models trained with 1–8 identities. Right, the same comparison for texture-based tasks. Blue line indicates CTFR-trained models (different identities with neutral expression); purple lines indicate CTFER models trained with different numbers of emotional variations (2, 4, 6, and 7 per identity). At any given number of training identities, increased emotional variation enhanced generalization in both geometry and texture domains more so than increasing identity variation, even when total training meshes (face categories) were matched (yellow markers used a similar total number of identity x emotion categories, n=7 or 8). **c-d,** Same as Fig. 4a**,b** but using a self-supervised pretrained Barlow Twins network as the base model for fine-tuning with colored-textured faces.

This synergy was seen as a consistent performance boost from emotion training across a range of identity set sizes. In the combined titration analysis, models trained with greater emotional variation consistently outperformed models with the same number of identities but less emotional variation, across both geometry- and texture-based tasks (**Fig. 4B**). Critically, emotional variation, which specifically affects face geometry, exerted a stronger effect on generalization than identity variation when models were matched for the total number of training categories (**Fig. 4B**; yellow markers). The training advantage from emotion was most apparent for geometry-based tasks, highlighting that the benefit of emotion-based training cannot be simply explained by mesh training set size alone (**Fig. 4B**; right).

### Emotion recognition training in textured faces improved performance across both supervised and self-supervised model pretraining

Thus far, we have examined how different face-training objectives influenced generalization across tasks. We next asked whether these effects depended on model initialization – specifically, whether our training approach is equally effective when applied to self-supervised versus supervised pretrained representations. We noticed that without any face fine-tuning, the ImageNet-pretrained self-supervised representation (Barlow Twins; BT-ResNet-50)^27^ already outperformed the supervised ResNet-50 baseline (Sup-ResNet-50) by a significant margin, achieving roughly 10% higher accuracy on CTFR and about 5% higher on GFR. This suggested that self-supervised pretraining provided more robust face representations off-the-shelf, even before domain-specific fine-tuning.

The stronger baseline of BT-ResNet-50 over Sup-ResNet-50 raises two possibilities: either further emotion-based fine-tuning would yield limited additional gains, or self-supervised initialization would synergize with supervised fine-tuning, as seen in prior semi-supervised settings^28^. Supporting the latter, fine-tuning BT-ResNet-50 with the CTFER-4IDx7EM face task led to consistent performance improvements across all face tasks. Because of the combined benefit of better off-the-shelf performance and further fine-tuning gains, the resulting model (BT-CTFER-4IDx7EM) outperformed its supervised counterpart (CTFER-4IDx7EM), with performance improvements of up to 8% on geometry-based face recognition (**Fig. 4C**). BT-CTFER models exhibited balanced gains across both geometry-and texture-based tasks, mirroring the generalization pattern originally observed with the supervised backbone but at a higher overall level of performance (compare wider area encompassed by model polygons in **Fig. 4C** vs. **4B**).

The combined emotion and identity titration analysis for the Barlow Twins base models revealed the same pattern observed for supervised ResNet-50 base models. Models trained with greater emotional variation consistently outperformed models with the same identity count but less emotional variation (**Fig. 4D**, blue vs. purple curves). As before, this advantage was most pronounced for geometry-based tasks when models were matched for the total number of training categories (**Fig. 4D**, yellow markers). Together, these results demonstrate that self-supervised pretraining synergizes with emotion-based face training, amplifying generalization and overall performance across face tasks.

### Face identity and emotion trained models learned representations that better preserved ground truth face distances as measured in 3D mesh geometry space

We next probed the nature of representations following face learning. If identity and emotion trained models extracted information about face geometry, we reasoned that this should be reflected in how representational distances between faces in model feature space aligned with ground-truth distances between the corresponding physical 3D face meshes. Because the vertices of all our faces were co-registered, we defined a geometric face mesh space based on the 936 vertices by 3 coordinates per vertex (2,808 dimensions). **Figure 5A** uses a low-dimensional PCA embedding to visualize the organization of 84 faces (12 identities x 7 emotions) in the physical mesh space (top 3 PCs, explaining 65.4% of total variance). One key insight was that the dominant axes of identity and emotion variation were approximately orthogonal: the angle between the corresponding vectors was 85.1° in the full PC space, indicating identity and emotion principally varied along approximately orthogonal directions in face space (**Fig. 5A**, red and blue arrows). Consistent with this structure, variance decomposition across principal components revealed that identity- and emotion-related variance loaded onto different PCs, with limited overlap between the components contributing most strongly to each factor (**Fig. 5B**).

**Figure 5.**
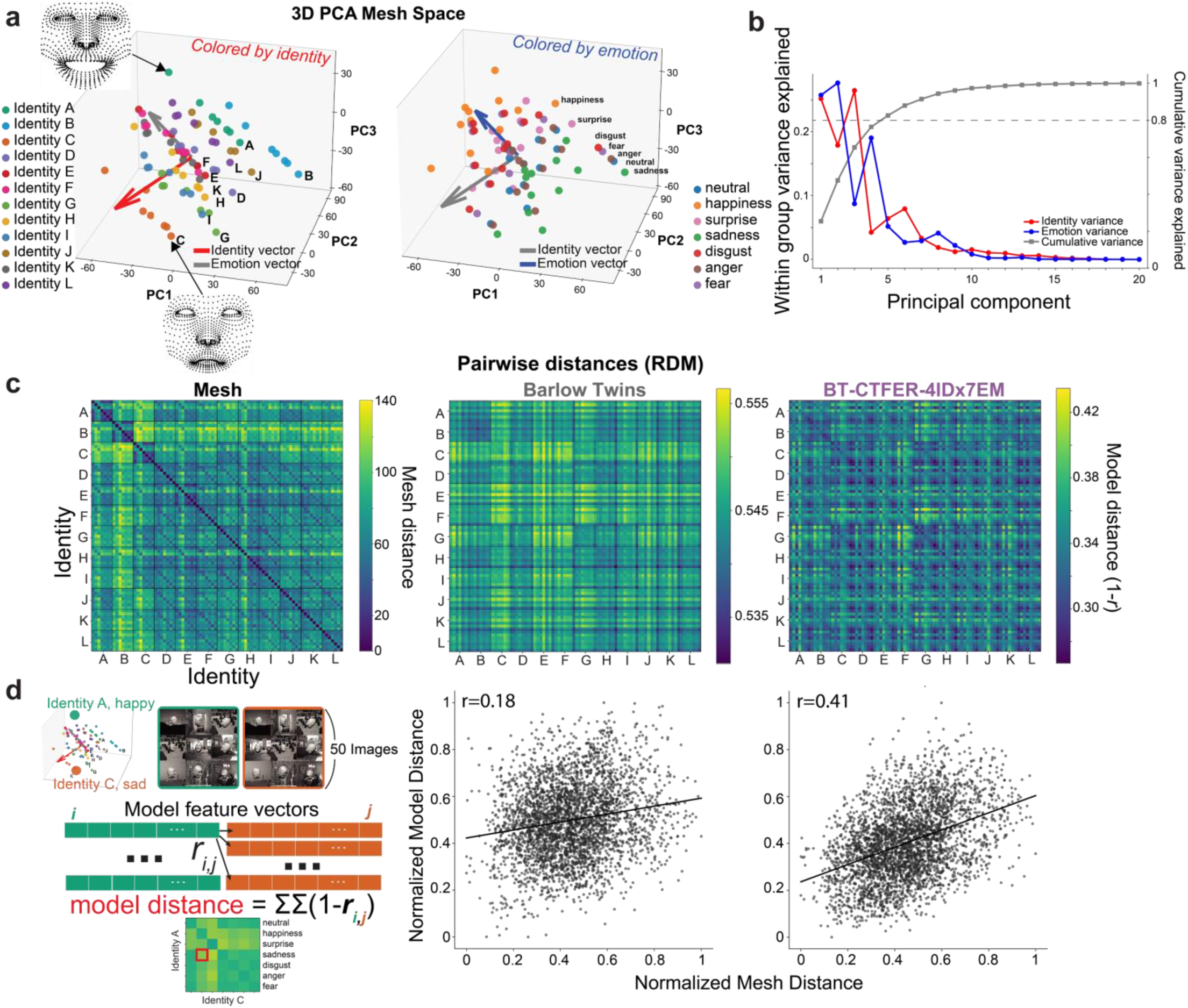
Representational distances between faces in the ground truth 3D mesh space compared to the learned feature space of the identity-emotion trained model. **a,** Visualization of the distances between face identities (left) and emotions (right) based on the ground truth physical mesh distances. Face space was constructed from the top three principal components (PCs) capturing variation across 84 face meshes (12 identities x 7 emotions). Identity and emotion variation engaged different directions of maximal variation in this space (red and blue arrows show directions of maximal identity-related and emotion-related variance, respectively). Two example face meshes at opposite extrema of these axes are shown (Face A_happy and Face C_sad). **b,** Relative contribution of each PC to identity (red) and emotion (blue) variance as well as to total variance (gray).The first five PCs accounted for 80% of the variance and generally contributed strongly to both identity and emotion variation but in contrasting proportions. **c,** Comparison of mesh-based and model-based representational dissimilarity matrices (RDMs). From left to right, RDMs based on: mesh vertex coordinates, self-supervised Barlow Twins ResNet-50 features, and features from the best-performing model (BT-CTFER-4IDx7EM). The BT-CTFER model captures the paradiagonal stripes driven by similarity of emotions across identities and evident in the mesh space RDM at left. **d,** Similarity between mesh space and model RDMs. Pearson correlation to the mesh space distances was higher for the BT-CTFER-4IDx7EM model than for the base version of Barlow Twins. Each dot in the scatter plot represents one pairwise distance between two faces, which for the models was computed by averaging distances across all pairs of images (n=50 image per mesh) from those two face meshes (illustrated at left using same example mesh pair as in Fig. 5a).

We then asked whether model representations preserved this geometric structure inherent to the face space defined by mesh distances. For this, we computed representational dissimilarity matrices (RDMs) from penultimate-layer activations of each trained model, averaging their representation of each of the 84 faces across 50 images drawn to yield 84 x 84-sized category-level RDMs.The RDMs were then compared to the ground-truth distance matrix via Spearman correlation. Qualitatively, the BT-CTFER-4IDx7EM model exhibited diagonal banding that mirrored the salient structure in the mesh-based distance matrix, a pattern that was largely absent from the RDM of the baseline Barlow Twins model (**Fig. 5C**). Quantitative analysis confirmed this difference, as pairwise distances between faces in the CTFER-trained model showed substantially stronger correspondence with distances in mesh-based PCA space than those of the baseline Barlow Twins model (**Fig. 5D**; *r_BT_*= 0.18, *r_BT-CTFER-4IDx7EM_*= 0.41). These results indicate that emotion-infused face training promotes representations that better recover the underlying geometric organization of face shapes, though there is only partial overlap with the ground truth mesh distance suggesting room for future improvement.

### Identity and emotion trained models improved match to human face recognition behavior

Besides representing faces in a manner more consistent with the underlying mesh space, we next asked whether these representations also produced humanlike face discrimination behavior. The idea that infusing rich emotion recognition can support robust face learning is consistent with how early human visual development likely unfolds. During infancy, a child spends countless hours closely interacting with a small set of familiar faces – parents and caregivers – who express a wide range of emotions. Under this hypothesis, we can ask whether we find an emergent phenomenon whereby models trained jointly on identity and emotion tend to produce humanlike patterns of face recognition behavior.

To quantify the match between models and humans, we conducted human psychophysics experiments to measure identity and emotion confusion matrices as behavioral proxies for perceptual similarity across different identity and expression categories (**Fig. 6B**, left). Participants performed a challenging two-alternative forced choice, match-to-sample face identification (7 identities) or emotion recognition (7 emotions), choosing which of two options matched a briefly flashed (200 ms) sample image. Humans performed both tasks at comparable accuracy levels (identity: 82%, emotion: 80%), and our leading model’s performance compared favorably (BT-CTFER-4IDx7EM, identity: 75%, emotion: 0.77).

**Figure 6.**
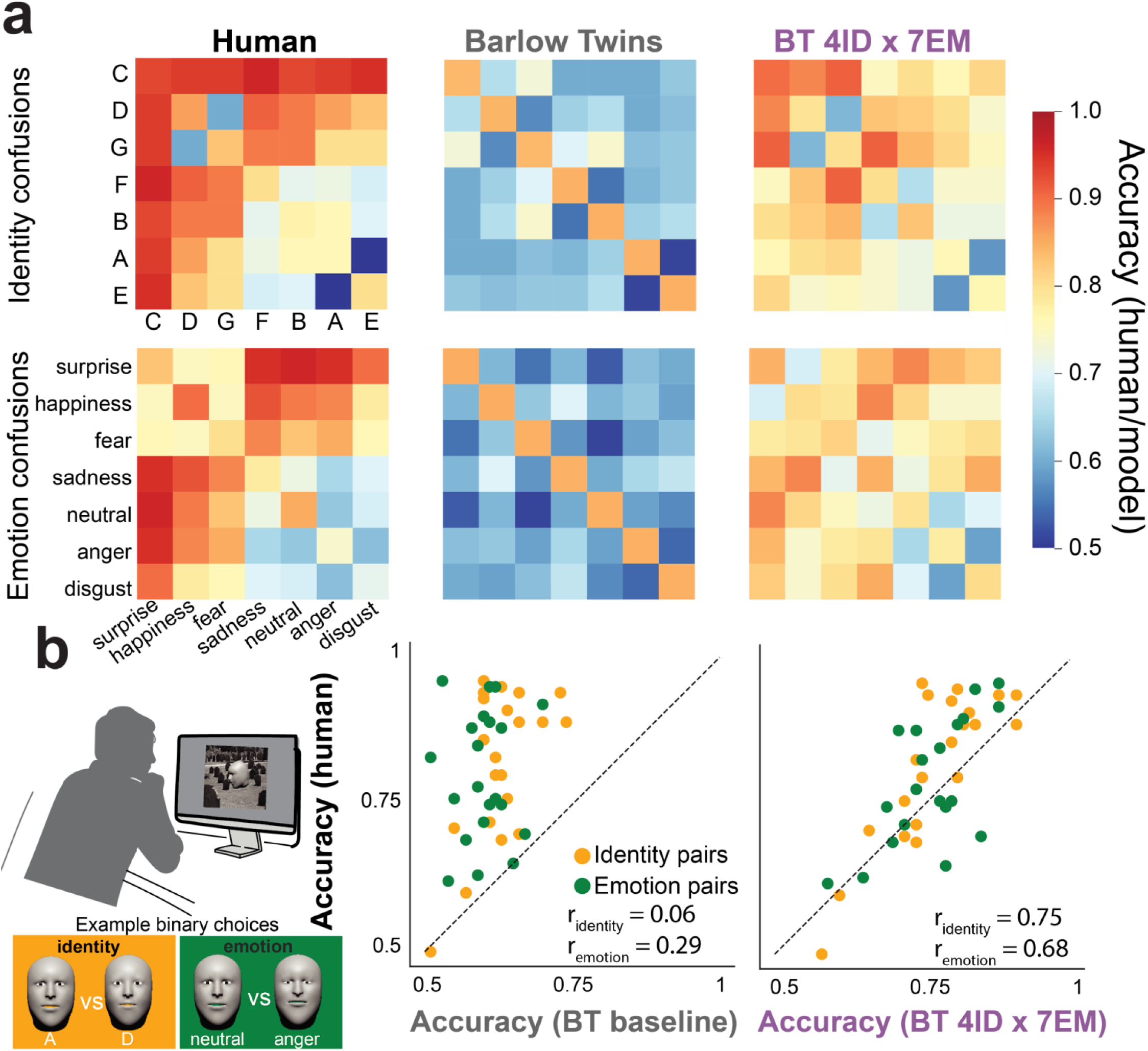
Correspondence between human and model behavior in face identity and emotion recognition. **a,** Pairwise discrimination accuracy matrices for identity (top) and emotion (bottom) tasks in humans (n = 5 subjects for identity, n = 6 for emotion; left) and the Barlow Twins and BT-CTFER-4IDx7EM models (middle and right). Each cell indicates mean accuracy for discriminating between a pair of identities (or emotions) when tested in a forced choice binary match-to-sample task. For visualization, mesh categories (rows) are sorted by human behavioral accuracy in descending order (high to low). **b,** Human and model confusion matrices for identity and emotion tasks (orange and green markers, respectively; each marker represents one pair of identities or one pair of emotions, see examples at left). The BT-CTFER-4IDx7EM model (right) exhibited classification errors that were much more humanlike than those in the base version of Barlow Twins (middle).

For humans, identity and emotion behavioral confusions were estimated with high reliability by pooling thousands of trials collected across multiple subjects (*r_identity, trial splits_* = 0.95, *r_emotion, trial splits_* = 0.95; n = 5 for identity, n = 6 for emotion) suggesting a robust human behavioral signature for comparing to models (**Fig. 6A**, left). The baseline ImageNet-pretained Barlow Twins model showed weak correlation with human confusions (BT-ResNet-50: *r_identity_* = 0.06, *r_emotion_* = 0.29). In contrast, our leading model that was trained with emotion variation across only four identities closely matched human identity confusion patterns and emotion confusion patterns (BT-CTFER-4IDx7EM: *r_identity_* = 0.75, *r_emotion_* = 0.68; **Fig. 6B**). Together, these results suggest that emotion-infused training improves not only overall performance but also fine-grained alignment with human perceptual judgments.

## Discussion

Face recognition remains a persistent challenge for artificial vision systems, despite being a domain where humans excel^29–31^. Our findings reveal a form of face representation that existing approaches cannot capture. Previous models trained on object recognition^11,15,32–34^ had shown that a form of face selectivity and identification performance can emerge from general object training without face-specific training. However, these successes relied on training and evaluation using static photographs that tend to emphasize texture and appearance cues and hence admit a wider space of recognition solutions. Our synthetic face framework took a different approach more focussed on geometry performance. Models could then be trained within a continuously parameterized 3D face space that systematically varied identity and expression information. This structured, natural shape information more closely mirrors real-world face transformations. By assessing generalization across identity versus emotion tasks and geometry versus texture conditions, this framework could reveal whether model encodings approached what might be expected from the invariant geometric relationships rather than from the surface-level 2D image features. In doing so, we shifted the criterion for biological plausibility from the mere presence of face selective model units and general face classification behavior to the capacity for geometry-based generalization, providing a deeper, more rigorous test of humanlike face processing.

Standard training regimes for expanding the face recognition abilities of discriminative models typically emphasize a large number of labeled identities when training with natural images containing faces, yet as we showed, this approach, even in the limit of a large number of experienced identities, likely fails to develop the geometric sensitivity that would lead to robust, generalizable facial understanding as found in biological vision^21,22,25^. Specifically, training on colored, textured faces failed to transfer performance to tests using non-textured, purely geometry defined faces (**Figure 4B**). Instead, transfer was much improved in the other direction, when training on geometry information alone in non-textured faces and then testing transfer to textured faces. However, since texture (e.g., hair, skin, and makeup) cannot be avoided in real world settings, we sought a more natural way to train networks than using textureless, purely geometry defined faces. Learning driven by emotion-based facial variations – within the same identity – produced representations with diverse benefits not strictly restricted to emotion recognition. These models developed geometric sensitivity that generalized effectively across facial appearances (texture to no texture) and diverse face tasks (emotion and identity) (**Figure 4**). In infusing emotion training, performance was even boosted on identity tasks compared to simply training on identity alone, and the learned representations on emotion and identity trained models improved the match to ground truth physical mesh space in their internal representations and best matched human confusion patterns in their output classification behavior.

Within the discriminative model framework, avoiding texture bias has long been a major Achilles’ heel to patch, and extensive prior work has considered input driven methods for avoiding texture biases in model training. These methods include low pass spatial filtering of images to suppress high spatial frequency information where texture information is concentrated^23,35–38^ and texture restylization approaches that replace original textures with synthetic ones, thereby making the shape the most diagnostic cue^22,39^. Although such inductive biases in the training diet can build some invariance to texture, we found that their effects on our specialized face task setting were limited and substantially weaker than the gains obtained from training on emotion-driven within-identity variation. Two factors may explain this discrepancy. First, our datasets included substantial 3D pose and lighting variation, which likely challenges models that rely on 2D shape cues, particularly those optimized on near frontal faces. Second, texture-debiased ImageNet models, trained for general object classification, may still lack the additional degree of fine-grained geometric sensitivity required for face recognition. Ultimately, we would venture that the visual system utilizes multiple mechanisms for internalizing geometric structure. Early in human development, limited spatial acuity and immature color vision would naturally downweight certain cues. For example, blurred, low color visual input would likely disfavor detailed texture information and perhaps robustify learning of face geometry information early on before the eyes and early visual machinery fully mature^23,40^. An additional intriguing biological learning mechanism early in development may arise from synergetic inductive biases in the motor behavior of caregivers: parents naturally bring their faces close to infants and produce exaggerated expressions, effectively tilting the infant’s input distribution toward high-signal geometric variation and reinforcing face-geometry learning^41–43^.

Thinking beyond initial face learning, where we posit that immutable geometry information provides a strong task-general foundation, room is still left for further precise refinement by fully leveraging identity-specific texture information, which likely supports peak identification performance as face expertise develops in adulthood. Consistent with this view, we did not observe the typical task tradeoff whereby training on one domain (geometry-defined faces) impaired performance in another (colored, textured faces). Rather, in our learning framework, geometry and texture task relevant features could be learned in parallel with minimal interference in a way that enabled broad generalization across all task types that we evaluated. This stands in contrast to control models trained on conventional unimodal classification using face photographs, which appeared to overfit in the domain of color face photographs and transferred poorly to other domains. These results suggest that geometry and texture likely operate in tandem in human perception, each conferring unique task-specific benefits. We therefore do not discount the parallel role of texture and appearance cues in face recognition, which has been extensively documented in the behavioral and neural literature^3,44,45^.

Having focussed here on enriching the training diet in the discriminative model framework for illustrating the effects of internalizing face geometry, it may yet prove equally sufficient to invoke explicit 3D objectives or 3D representations to further encourage face geometry learning. However, this may not be strictly necessary since geometry-rich emotion signals, like those we used to train our models, are already abundant in natural human experience. Expressive, exaggerated parental interactions during infancy^46^ provide high-geometry, low-texture-biased visual input, and this supply only increases as social interactions become more varied and nuanced throughout development. Emotion or further articulation learning therefore remains a strong and biologically plausible candidate mechanism for promoting geometry-based face representations even when staying within the architectures and objectives of the standard discriminative model framework. Future work is needed to explore whether emotion learning in explicitly 3D models forms a potent combination that further augments face recognition performance.

It’s worth noting that emotion-based face learning proved to be quite efficient. First, without even tilting the low-level statistics of training images (e.g., removing confounding texture) or introducing any explicit 3D priors into the loss function or model architecture, models trained solely on emotion variation generalized robustly to geometry-based tasks. Second, emotion-based training improved sample efficiency by yielding multiple training images from just a single identity. Arguably, a single face can generate a seemingly unlimited number of unique expressions, potentially on the same order as the number of distinct identities one might encounter. Empirically, we found that without any identity variation, emotion variation from a single individual was sufficient for models to learn to identify new faces, impressively generalizing their performance from emotion recognition over to identity recognition tasks. Rather than be narrow in their variation and hence limited in training utility, emotion-based deformations provided meaningful geometry information. This was reflected in consistent performance improvements as the number of expressions increased from two to seven during training. Future work can determine how much a rich, densely sampled space of expression intensities (e.g., half smiles versus full smile), other expression types (e.g., confusion, contempt), and additional articulations and gestures (e.g., vocalizing, winking) beyond the seven basic emotions that we used can further aid model performance. In the future, an additional route for gaining efficiency would be to marry emotion variation with the power of self-supervised training methods, thereby reducing or potentially eliminating the need for discriminative face labels in early training. Training could proceed from unlabeled natural movies of human interaction where moment-to-moment changes in facial geometry such as a speaker’s mouth and facial movements or a listener’s emotional reactions create natural, identity-preserving 3D transforms that could serve as positive or negative pairs for self-supervised contrastive learning.

In the realm of human psychology and behavior, this newfound perspective may help explain what makes human face processing computationally special relative to general object recognition: humans develop into face articulation experts, specialized in decoding fine-grained deformations of geometric structure. Here, our work suggests that leveraging natural emotional variation in early human experience may lead to computationally critical benefits and suggests a new, more biologically inspired, efficient process for visual learning in artificial vision systems.

## Acknowledgements

This work is supported by the funds provided by the National Science Foundation and by DoD OUSD (R&E) under Cooperative Agreement PHY-2229929 (The NSF AI Institute for Artificial and Natural Intelligence), Klingenstein-Simons fellowship, Sloan Foundation fellowship, and Grossman-Kavli Scholar Award. This work was performed on the Columbia Zuckerman Institute Axon GPU cluster.

## Author Contributions

**Seojin Lee**, Conceptualization & Analysis – modeling experiments, Writing – original draft, and Writing – review and editing

**Josh Ying**, Conceptualization & Analysis – early modeling experiments

**Ahana Dey**, Conceptualization & Analysis – behavioral experiments

**You-Nah Jeon**, Software, Stimulus design, Conceptualization – early behavioral experiments

**Elias B. Issa**, Conceptualization – all stages of modeling & behavior experiments, Resources, Supervision, Funding Acquisition, Methodology, Writing – original draft, Project Administration, Writing – review and editing

## Methods

### Face stimuli

#### Capturing faces

We curated a dataset of twelve identities for the Geometric Face Recognition (GFR) task using a custom iPhone application based on Apple’s ARKit API (https://developer.apple.com/documentation/arkit/arkit_in_ios/content_anchors/tracking_and_visualizing_faces). ARKit accesses the mobile device’s built-in TrueDepth camera, which projects an infrared pattern of tens of thousands of dots onto the face which effectively serve as fiducial markers for reconstructing a dense 3D depth map in realtime. The data is processed by ARKit to generate an ARFaceAnchor object, which outputs the 1220 vertex coordinates describing the 3D face mesh and 52 BlendShape coefficients that encode the magnitude of individual facial movements to extract 3D mesh coordinates, BlendShape coefficients, the projective transform matrix, and the corresponding 2D image frames^47^. Videos were recorded on an iPhone 14 under consistent lighting and background conditions. Participants (n = 12) spanned a wide range of ages, genders, and racial and ethnic backgrounds. Each recording produced a text file containing 1220 x 3 vertex coordinates and 52 BlendShape coefficients per frame. For each identity, a single frame displaying a neutral facial expression was selected as the base mesh.

#### Stitching faces

To generate composite 3D faces, we stitched the collected frontal face mesh onto a base faceless 3D dummy head using custom MATLAB code (https://github.com/issalab/Jeon-apple_facestim_generation). From each mesh, the identical 936 centrally located vertices were selected to isolate the internal facial structure. The mesh was then aligned to the base head by matching vertex normals at the attachment point, and boundaries were smoothed using a distance-weighted averaging method. To each fused face-head mesh, identical eyes (i.e., spheres for pupil and sclera) were added in Blender (www.blender.org), and the final mesh was smoothed using the *Smooth Faces* function in Blender. The base head and eye meshes were kept identical across identities to ensure that differences reflected the internal face geometry alone.

#### Generating face training datasets

From each identity, we rendered 50k training images with varying 3D viewing parameters (size, rotation, and lighting) using MkTurk (mkturk.com)^48^, a web-based behavioral platform for realtime 3D rendering and psychophysical experiments developed in the lab (code available at https://github.com/issalab/mkturk). Each 3D face was placed centrally on a grayscale natural-image background randomly selected from 1,428 ADE20K images (https://ade20k.csail.mit.edu/index.html). Face size was uniformly sampled from a uniform distribution (0.550 to 1.155) in Three.js 3D world units, relative to a 3.0925-unit sized canvas. All rendered images were exported at 461 × 461 pixels and resized to 224 × 224 pixels before being passed to the model. Horizontal rotation varied between −90° and +90°, and vertical rotation between −45° and +45° (with visibility constraints). Lighting position was drawn uniformly from (*x*, *y* ∈ [−1, 1]; *z* = 1, where *x* is azimuth and *y* is elevation, and the light always pointed at the origin [0,0,0]).

Model behavioral testing sets contained 151 images per identity generated using the same rendering pipeline but with distinct parameters. Ten background scenes were each sampled 15 times (15 images using the same one of the ten backgrounds). Face size was drawn from 0.37–0.765 (Three.js 3D world units; canvas = 1.61 units) to match previous behavioral datasets used in the lab. Testing images included upright (0°) and inverted (+180°) views and the three lighting contexts: overhead downlighting, underneath uplighting, and contrast reversal (both face and background). Combining the upright/inverted conditions with the three lighting contexts yielded six conditions total per identity (25 images in each condition). An additional upright, front-view image was rendered without a background for reference (face token image).

#### Generating textured face stimuli

To create textured meshes, we projected the 2D frontal-view photographs from FaceCaptureX onto each corresponding 3D face. Each photo was manually edited in Adobe Photoshop to remove non-facial regions and smoothed to minimize high-frequency detail. UV coordinates (texture coordinates) were computed in MATLAB by mapping mesh vertices through world-to-camera transformations, overlaid to confirm alignment, and extended with skin-tone fill to cover all UV points. The final textured mesh was generated using GLTF functions for applying the edited photo as a texture map. Textured meshes were rendered using the same MkTurk image-generation pipeline, except that the grayscaled ADE20K backgrounds were replaced by colorized ADE20K images.

#### Generating emotion face stimuli

The BlendShape representation of Apple ARKit encodes facial expressions using 52 predefined coefficients weighting a direction of change defined by some combination of face mesh vertices. Each recording provided co-registered 3D vertices (1220 x 3) and BlendShape vectors per frame. To estimate how each BlendShape affects vertex displacement, we fit a ridge-regression model Y = XW with penalty = 0.1, where *X* ∈ *R*^*n*×53^ (52 coefficients + bias) and *Y* ∈ *R*^*n*×3660^ (flattened 3D coordinates). The resulting matrix *W* ∈ *R*^53×3660^ describes vertex-wise influence for each coefficient. We defined six target emotions – anger, disgust, fear, happiness, sadness, and surprise – plus the neutral condition, creating seven expression states. Each emotion was represented by a manually specified 52-dimensional BlendShape vector that approximated target frames where a reference subject was instructed to create that expression. The corresponding deformation was computed for *n* frames. We then performed a ridge regression to estimate the linear mapping: *ΔY = X^T^W*, reshaped into 1220 x 3 and added this deformation to the neutral mesh. Because *W* encodes identity-independent deformation for vertices that are co-registered across individuals, any neutral identity could be converted to reflect each emotional expression. Expression intensity was normalized by total vertex displacement (sum of Euclidean distances between neutral and emotion meshes). Displacements ranged from 3.27–7.36 units; each was rescaled to a reference intensity of 4 to ensure natural appearance. These normalized meshes were rendered into training and testing images using our 3D rendering pipeline.

#### Basel Face Stimuli (texture and no texture)

The Basel Face Model (BFM; https://faces.dmi.unibas.ch/bfm/index.php?nav=1-1-0&id=details) is a publicly available 3D Morphable Face Model built from high-resolution 3D scans of 100 female and 100 male individuals, representing each face with 53,490 co-registered vertices^24^. Shape and texture are modeled by independent principal components in vertex space. To parallel the GFR and CTFR image datasets created from our in-house Apple-based face mesh stimuli, we selected twelve identities from the BFM to match the number of individuals used in the Apple-based stimuli. Each identity was rendered in two forms: (1) textured, using the BFM albedo, and (2) shape-only, with uniform surface shading. All stimuli were rendered with the same image-generation pipeline used for the Apple-based faces.

#### Face Photographs

A two-identity photographic dataset was constructed from a subset of the *Labeled Faces of the Wild (LFW)* corpus^49^. Two high-variability identities, George Bush and Collin Powell, were selected. From each, 151 images were randomly sampled (302 total). These photographs contained natural variation of the face’s viewpoint, lighting, expression, and background.

### Tasks

#### Geometric face recognition tasks

We evaluated the DNN models on two geometry-based face recognition tasks: **Geometric Face Recognition (GFR)** and **Geometric Emotion Recognition (GER)**.

**GFR** comprised 66 binary discrimination tasks, each involving all possible pairs of identities selected from a pool of 12 unique individuals. The design paralleled the two-alternative forced choice (2AFC) face discrimination paradigm used in our prior behavioral work, enabling direct comparison with human and monkey performance in a two-way (binary) face identity discrimination task. On each trial, feature activations were extracted from the model for images of two identities, and a linear classifier was trained to predict image identity. Recognition accuracy for each identity pair was computed at a given model layer, and mean accuracy across all 66 pairs was used as a measure of model performance.

**GER** assessed within-identity discrimination by requiring models to distinguish between emotional expressions of the same individual (e.g., *neutral* vs. *happiness* in Identity A). The task included 21 binary discriminations spanning all pairwise combinations of seven emotions. To simulate generalization of emotion discrimination to unseen identities, one identity that was held out during training was used for evaluation.

#### Texture-augmented face recognition tasks

To assess the generalization of geometry-trained models to texture-rich domains, we curated **Color-Textured Face Recognition (CTFR)** and **Color-Textured Emotion Recognition (CTER)** – texture-augmented analogues of images used in GFR and GER. These tasks used identical identity and emotion combinations but included colorized facial textures and backgrounds. CTFR and CTER therefore tested whether models trained on geometry-based variation would generalize to domains with superficial appearance and texture cues.

#### Basel face recognition tasks

We further tested model generalization using the Basel Face Model (see *Stimuli*). Twelve identities were selected from the Basel dataset to match the number of individuals used in the GFR and CTFR tasks. Each task – **Basel-GFR** (shape-only) and **Basel-CTFR** (textured) – consisted of 66 binary discriminations, spanning all pairwise combinations of the twelve Basel identities. These tasks evaluated whether geometry-based training enabled generalization to a qualitatively different set of novel 3D morphable faces from the Basel framework that differed in identity as well as higher geometric mesh resolution from the training set (53,490 versus 1,220 vertices).

#### Natural face image control task

The **Face Photograph** task evaluated the model performance on real face images. Two identities were selected from the Labeled Faces in the Wild (LFW) dataset, each with 151 unconstrained photographs. This task served as a naturalistic face recognition benchmark against which synthetic face performance could be compared.

### Computational models

#### Model architecture

We leveraged deep neural networks (DNNs) as computational models to simulate feature extraction, high-dimensional representation, and recognition processes. The primary architecture used in the study was a ResNet-50^18^ backbone pretrained with supervised learning on the ImageNet dataset, a widely used benchmark for large-scale object classification. To assess the generality of our findings across architectures, we additionally evaluated two baseline models: AlexNet^19^ and VGG-16^20^, both pretrained on ImageNet. In addition to object-trained models, we included two architectures specialized for face recognition: VGG-Face^8^, a VGG-16-based model trained to classify 2,622 celebrity identities using a large-scale face dataset (∼2.6M images), and two FaceNet^7^ variants, both implemented using the InceptionResNetV1 architecture from the facenet_pytorch library. One variant (FaceNet-128D) was pretrained on the CASIA-WebFace dataset and produces a 128-dimensional L2-normalized embedding, closely following the original FaceNet formulation^7^, which used a triplet loss to directly optimize the embedding for face similarity. The other variant (FaceNet-512D) was pretrained on VGGFace2 and produces a 512-dimensional embedding. These pretrained models (FaceNet-128D and FaceNet-512D), while they use classification-based objectives rather than triplet loss, retained the compact embedding design and identity discrimination focus of FaceNet. These face-specialized architectures thus served as reference networks for specifically optimizing for identity discrimination in real-life face photographs.

In developing new face training methods, we further explored the influence of our training objectives and training diet during fine-tuning on previously learned image representations by testing with supervised trained ResNet-50 as well as a variant of ResNet-50 pretrained with the Barlow Twins contrastive learning objective^27^ on ImageNet. Unlike standard supervised training, Barlow Twins employs a self-supervised objective that encourages invariance in the latent variable space in concert with redundancy reduction between two different augmented views of the same image, enabling the network to organize object and scene representations without relying on explicit class labels. By including this model, we tested whether less supervised, representational structures emerging from ImageNet pretraining better supported geometry-based recognition tasks.

#### Fine-tuning

We fine-tuned ResNet-50 models pretrained on ImageNet classification or Barlow Twins self-supervised learning objectives. For fine-tuning an ImageNet-pretrained ResNet-50, the final fully connected (fc) layer was removed and replaced with a randomly initialized linear classifier matching the number of target identity classes. The rest of the backbone, including all convolutional blocks and the global average pooling layer, was retained and fine-tuned end-to-end. For Barlow Twins-pretrained ResNet-50, the original projection head (a 3-layer MLP after the average pooling layer) was removed, and a new randomly initialized linear classification layer was appended.

Both model types were optimized using stochastic gradient descent (SGD) with a learning rate of 0.001, momentum of 0.9, and weight decay of 1e-4. The learning rate was decayed by a factor of 10 every 500 steps using a StepLR scheduler. Fine-tuning was conducted for 2 epochs with a batch size of 64. The training set contained 100,000 images in total, split 80/20 between the training and validation subsets. Data augmentation included random resized crops and horizontal flips, followed by ImageNet normalization. Model checkpoints were saved every epoch, and the best-performing checkpoint based on validation accuracy was used for downstream analyses.

#### Face pair selection for fine-tuning

To ensure fair comparison with human and monkey behavioral data, we held out the two identities A and B from model training that were used in those behavioral data collections, and used these identities for model evaluation only. Among the remaining 10 identities C-L, we selected subsets of face pairs with increasing number of identities included: three 2-identity subsets (C and D, C and E, D and F), followed by one 4-identity set (C, D, E, F), one 6-identity set (C, D, E, F, G, H), and one 8-identity set (C, D, E, F, G, H, I, J). For the 2-identity condition, we deliberately used multiple identity pairs to reduce the chance that results were driven by a specific face pairing. For larger identity sets, the risk of such bias was lower, so a single configuration per set size was sufficient. We limited the largest set to 8 identities to leave four identities unseen during training, enabling more comprehensive out-of-distribution (OOD) evaluation. Following the initial training of shape-only stimuli (GFR), we trained models on the corresponding textured faces (CTFR) using the exact same identity subsets.

We next introduced emotional variation (CTFER), expanding the training categories by varying facial expressions. For identity-emotion-mixed training (IDEM), we varied both the number of identities and the number of emotional expressions per identity. In the main setup, we trained models on two identities (C, D) across all seven emotional expressions (neutral, happiness, sadness, fear, anger, surprise, disgust) to form a 14-way IDEM task (each identity x emotion pair formed a class). This was expanded to a 28-way IDEM task by using four identities (C, D, E, F), each across the same seven emotions. We additionally tested training with a subset of emotions (e.g., neutral and happiness only) to assess the impact of emotional diversity on model performance.

#### Model layer selection

Across different architectures, we extracted model representations from the penultimate layer – the final stage before classification heads in supervised models (e.g., supervised ResNet-50, AlexNet, VGG-16) and before the embedding head in contrastive models (e.g., Barlow Twins ResNet-50). This stage typically corresponded to high-level, abstract visual processing, capturing shape- and identity-related information without committing to explicit class outputs driven by the original learning objective. For FaceNet, we similarly used the final 128-dimensional embedding. Overall, the layer choice ensures comparability across models and mirrors prior work showing that the penultimate stage best aligns with behavioral and neural representations^447,49–51^ Specifically, we extracted features from the final average pooling layer (avgpool) of the supervised ResNet-50 and Barlow Twins ResNet-50 models, the final fully connected layer (fc7) of VGG-16 and VGG-Face, the penultimate fully connected layer (fc7) of AlexNet, and the standardized 128-dimensional embedding layer of FaceNet.

#### Linear classifier readout of model face discrimination

To evaluate a given model’s face discrimination behavior, we simulated a 2AFC task between pairs of categories using binary classifier readouts (i.e., different pairs of faces). For each pairwise task, we extracted features from all images in the two categories, trained a linear support vector machine (SVM) classifier (hinge loss, L2 penalty, C=1.0), and evaluated performance using two-fold cross-validation repeated 100 times with random splits of the training images. Each category pair was loaded from pre-defined image folder structures corresponding to its associated task (e.g., 66 identity pairs in GFR, 21 emotion pairs in GER). We evaluated classification accuracy separately for each pair and then averaged across pairs to obtain the overall dataset-level classification accuracy score. We report the mean classification accuracy and standard deviation across repetitions for each model and task. Additionally, we computed category level confusion matrices between meshes (each mesh reflected a particular identity x emotion), where performance on each mesh pair was computed by averaging discrimination accuracy across 20 rendered images per mesh.

### Representation analyses

#### Face space

To characterize the structure of identity and emotion variation in raw face geometry, we performed a principal component analysis (PCA) directly on 3D face meshes, specifically the matrix of vertex coordinates. The goal was to obtain a low-dimensional “face mesh space” capturing the main axes of shape variation across identities and emotional expressions. We used meshes collected from 12 individual identities across 7 emotional expressions (see **Methods** “Capturing Faces”), which resulted in 84 total identity-expression combinations. Each mesh was represented as a flattened vector of concatenated *x*, *y*, and *z* coordinates across 936 vertices (resulting in 2808 dimensions per mesh). The mesh coordinates were first centered by subtracting the mean shape across all 84 faces and then standardized by dividing by the per-dimension standard deviation. This normalization ensured that subsequent principal components captured shape variation independent of scale. PCA was performed on the standardized mesh vectors, yielding a set of orthogonal axes ranked by explained variance. The top three principal components (PC1: 25.41%, PC2: 22.09%, PC3: 17.87%) together explained 65.36% of the total variance and defined a low-dimensional face mesh space summarizing the main axes of geometric variation across identities and emotions.

#### Ground-truth distance matrix

To quantify geometric relationships among all faces, we computed Euclidean distances between every pair of meshes in the full PCA space, using all retained components derived from the standardized 3D mesh data. The resulting 84 x 84 face-distance matrix defined the ground-truth representational geometry of the stimulus set, capturing physical shape differences across identities and expressions. This matrix served as the reference structure for model-geometry comparisons in subsequent analyses.

#### Quantifying Identity and Emotion Axes

To assess whether identity and emotion varied along distinct directions in PCA-derived face space, we first computed a centroid for each identity (averaged across all emotions) and each emotion (averaged across identities). Covariance matrices were computed separately for these centroids, and the eigendecomposition of each yielded orthogonal axes representing the dominant directions of identity and emotion variation.

The top eigenvector for each covariance matrix defined the principal identity and emotion separation vectors. The angle between them was computed using the arccosine of their cosine similarity:

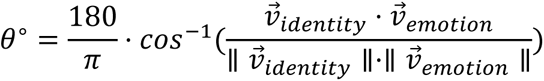

This yielded an angle of 94.87°, indicating near-orthogonality between identity and emotion dimensions. A similar relationship was observed within the reduced, low dimensional 3D PCA subspace, which was used for visualization in **Fig. 7A-B**.

#### Model representational dissimilarity matrices (RDMs)

To compare model representations to this geometric structure of face space derived from mesh coordinates, we computed representational dissimilarity matrices (RDMs) for 84 face categories (12 identities x 7 emotions). Each category contained 50 images drawn from the same 3D mesh rendered with randomized lighting, viewpoint, size, and background. Feature activations were extracted from the penultimate layer of each model for all 4,200 images (84 categories x 50 images). Pairwise dissimilarities were computed as 1 - Pearson’s correlation between feature vectors, yielding 4,200 x 4,200 image-level RDM. To obtain a category-level RDM, this matrix was divided into 84 x 84 blocks, each corresponding to a pair of identity-emotion categories. Averaging each 50 x 50 block yielded a single dissimilarity value per category, resulting in an 84 x 84 block-averaged RDM summarizing mean representational distances between categories.

#### Face-space-to-model-representation correlations

To quantify the correspondence between the ground-truth face space that captures the geometric relationship between faces and models’ representations, we correlated each model’s block-averaged RDM with the PCA-derived distance matrix. Both matrices were vectorized by extracting their upper-triangular elements, and their correspondence was quantified using Spearman’s rank correlation (ρ). This analysis assessed how well each model’s representational geometry preserved the true geometric relationships among faces.

### Behavioral Experiments

#### Prior human behavioral data

Human face discrimination performance in **Fig. 1** was collected in our prior work for a binary discrimination task between one specific pair of faces^17^. Briefly, each trial began when the subject touched a central fixation point, followed by a brief presentation of a sample image (200 ms duration). Two-choice images were then displayed, corresponding to the two possible face categories and with fixed choice positions across trials to maintain consistent stimulus-response mapping. Subjects selected the matching image by touch.

#### Human identity and emotion discrimination experiment

Here, we sought to extend our prior work to measure human performance across multiple target-distractor face pairs to generate a confusion matrix. These human behavioral data were collected in the laboratory using MkTurk. Two tasks were administered: a 7-way identity discrimination task and a 7-way emotion discrimination task, where emotion intensities were scaled such that the average vertex displacement matched that from identity changes in mesh space. This normalization in mesh space helped bring the identity and emotion discriminations closer in difficulty. A match-to-sample task structure was used where there were two possible choices, and the goal of the subject was to match the identity (or emotion) in the sample image to that in one of the two choices. The MtS design allowed us to probe binary tasks while expanding the sample category set to include seven identities or seven emotions, yielding 21 pairwise discriminations between all pairs of the seven meshes in the identity, or emotion task.

Each trial began with a fixation point at the center of the screen. Participants maintained fixation for 40 ms to initiate a trial, after which a sample image was presented for 200 ms. In the identity task, sample images were drawn from seven face identities under neutral expression. In the emotion task, images were drawn from seven emotion categories of a single individual — anger, disgust, fear, happiness, neutral, sadness, and surprise. For both tasks, each category contained 20 unique images, sampled uniformly with replacement from the corresponding image sets to ensure balanced category frequencies. Each trial allowed up to 5 seconds for a response, with no inter-trial interval to maintain continuous task flow. Participants completed 1,000–2,000 trials for each task. The identity task was completed by five participants, and the emotion task by six participants recruited from the Columbia University community. All participants had normal or corrected-to-normal vision and provided informed consent.

### Human and Model Behavioral Signatures

#### Model confusion matrices

For each model, we computed two identity- and emotion-based confusion matrices using stimuli from the colorless and textureless GFR and GER datasets. To parallel the behavioral experiments, evaluation was restricted to a subset of seven identities, which gave 7 x 7 matrices aligned with the category structure used in the behavioral experiments. Feature activations were extracted from the penultimate layer for all evaluation stimuli. To quantify discriminability, linear classifiers were fit in a one-vs-one manner across all 21 category pairs (seven identities or seven emotions), using the same 20 images per category that were used in the behavioral task. Classification performance was estimated using cross-validation on the 20 images per category, matching the behavioral tasks. Accuracies were averaged across folds and assembled into the full confusion matrix, with diagonal elements indicating correct classifications (O1) and off-diagonal elements indicating pairwise confusions (O2)^450^ Each matrix was row-normalized to allow comparison across categories.

#### Human confusion matrices

Behavioral confusion matrices were derived from the behavioral datasets described above, using the same subset of GFR and GER image sets analyzed in the models. For each subject and task, the probability of choosing category *j* when category *i* was presented was computed from trial-level choice data, yielding a 7 × 7 matrix normalized by row. Individual matrices were then averaged across subjects to produce a population-level behavioral matrix.

#### Human-model correlations

To quantify alignment between model and human performance, we vectorized the off-diagonal entries (O2) of each confusion matrix, which capture the pattern of pairwise category confusions. Spearman rank correlations (ρ) were computed between the model and human O2 vectors for each task. This provided a single measure of behavioral correspondence that reflects the structure of confusions beyond overall accuracy.

### Code Availability

Image generation and psychophysics code is available at https://github.com/issalab/mkturk.

